# Novel Laser Capture Microdissection-Proteomic Analysis Identifies Spatially Distinct Extracellular Matrix Signatures in the Core and Infiltrating Edge of Human Glioblastoma

**DOI:** 10.1101/2022.09.01.506199

**Authors:** Robert Pedley, Danielah T. Prescott, Ellen Appleton, Lewis Dingle, James Minshull, Pietro Ivo D’Urso, Ibrahim Djoukhadar, Andrew P. Gilmore, Federico Roncaroli, Joe Swift

## Abstract

**Background:** Glioblastoma is the most common and aggressive primary brain tumour in adults. Hallmarks of glioblastoma include its intra-tumoural heterogeneity and extensive invasion of the surrounding brain. Glioblastoma is known to remodel the extracellular matrix (ECM) of the brain, resulting in altered mechanical properties and the establishment of a tumour-promoting microenvironment. How changes in the expression and spatial distribution of ECM constituents within glioblastoma contribute to invasion and disease progression is still unclear.

**Methods:** Here we report on a protocol for laser-capture microdissection coupled with mass spectrometry (LCM-proteomics) that allowed a spatially resolved and unbiased analysis of the regional ECM proteome (matrisome) in formalin-fixed and paraffin-embedded (FFPE) samples of human glioblastoma. We investigated five molecularly characterised hemispheric adult glioblastomas where the brain/tumour interface and tumour epicentre were represented in the surgical specimens and snap-frozen tissue was available. LCM-proteomic analysis was validated with immunohistochemistry.

**Results:** LCM-proteomics identified 53 matrisome proteins in FFPE tissue, demonstrating comparable performance with conventional analysis of snap-frozen tissue. The analysis revealed distinct matrisome components in the brain/tumour interface versus the tumour epicentre. Guided by data from LCM-proteomic analysis, immunostaining for tenascin-R confirmed greater staining in the brain/tumour interface, whilst expression of fibronectin was higher in the tumour epicentre.

**Conclusion:** The protocol described in this work allowed for accurate, spatially resolved analysis of ECM in FFPE tissues, with performance comparable to analysis of snap-frozen tissue. While the focus for this work was on the regional ECM composition of glioblastoma, we found that the LCM-proteomics protocol is also applicable to the study of the wider proteome, and represents a valuable tool for investigating tumour/tissue heterogeneity. This protocol opens the possibility to apply LCM-proteomics to retrospective studies with the advantage of accessing clinical history and follow-up information, providing a valuable resource for translational research in glioblastoma.

## Introduction

Glioblastoma is the most common and aggressive primary malignant brain tumour in adults. Tumours of this type are characterized by extensive infiltration of the normal brain and marked intra- and inter-tumour molecular heterogeneity (1–3). Despite the progress of surgical technique and a more robust molecular stratification to guide chemo-radiotherapy, recurrence is the norm and median overall survival is 15-18 months from diagnosis (4–6).

The tumour microenvironment (TME) plays a complex and dynamic role in the pathobiology and progression of glioblastoma. The extracellular matrix (ECM) is a key constituent of the TME and regulates cellular proliferation and migration through biophysical and biochemical cues (7, 8). Although it is accepted that ECM structure and composition is altered in glioblastoma, the full spectrum of molecular changes still remain elusive. Neurosurgeons describe differences in the texture and density of tumour tissue when compared with normal brain parenchyma, which is consistent with the quantitative observation that ECM stiffness is increased in glioblastoma tissue relative to low-grade gliomas (9). Glioblastomas have been shown to express atypical ECM components, or express ECM at abnormal levels to that of the healthy brain parenchyma (10). Aberrantly regulated ECM components include: collagens (such as type IV, III, I) (11–13); glycoproteins (such as tenascin-C) (14–16); proteoglycans (e.g. versican & perlecan) (17, 18) and matrix metalloproteinases (such as MMP-1,2 &9) (19, 20). *In vitro* studies have shown that glioblastoma derived cells are sensitive to the presence of various ECM substrates (21–23). Furthermore, glioblastoma cells are responsive to the mechanical properties of their environments, exhibiting increased proliferation and migration on stiff substrates when compared to soft substrates (9, 24, 25).

Intra-tumour spatial heterogeneity of tumour cells and the TME influences clinical outcome (1–3). The epicentre (referred to henceforth as ‘core’) of glioblastoma is a hypoxic, pro-inflammatory and mechanically stiff environment. Multiple studies have documented the malignant effect of these conditions on neoplastic and non-neoplastic cells (9, 26, 27). In contrast, the tumour/brain interface (referred to henceforth as the ‘leading/infiltrating edge’), whilst appearing similar in structure and composition to normal brain tissue under microscopic examination, accounts for the location of 90% of recurrences that occur following surgical resection (28, 29). Recent evidence suggests that glioma stem cells migrate from the tumour core into the infiltrating edge, where they contribute to the remodelling of the tissue microenvironment to produce a tumour-promoting niche (29–31). However, despite the clinical importance of the tumour/brain boundary, relatively few studies have examined the microenvironment of the leading edge in detail. There is, therefore, a need for detailed studies that spatially resolve the glioblastoma TME in order to decipher the influence of regional microenvironments on glioblastoma pathobiology and disease progression.

Laser-capture microdissection (LCM) is a methodology that allows the excision and collection of discrete regions from tissue sections under microscopic visualisation (32). Isolated tissue can then be subjected to a variety of downstream analyses. Recently, LCM was utilised in combination with RNA-sequencing to study the expression profiles of distinct histological regions within glioblastoma (33), and with liquid chromatography tandem mass spectrometry (LC-MS/MS) to explore the ECM composition of the lung microanatomy (34). Herein, we couple LCM with LC-MS/MS to perform a spatially resolved investigation into the extracellular matrix biology of the core and leading edge of human glioblastoma. We demonstrate the feasibility of analysing the glioblastoma tumour microenvironment from archived FFPE tissue, and identify signature ECM compositions within these defined tumour compartments.

## Materials and Methods

### Ethical approval

The study has been ethically approved (REC reference: 19/NE/0186 IRAS project ID: 244538)

### Specimen selection

The tissue from five de novo IDH1/2 wild-type glioblastomas that underwent extensive surgical debulking were retrieved from the archives of the department of Cellular Pathology at the Northern Care Alliance, Salford Royal, United Kingdom. All specimens included the tumour / brain interface that could be orientated and compared to pre-operative neuroimaging. Genetic and epigenetic profile was assessed in all five cases. Matched snap-frozen tissue from the specimens were available.

### Laser capture micro-dissection (LCM) of formalin fixed paraffin embedded (FFPE) tissue

5μm sections of (FFPE) tissue were mounted onto either glass slides or MMI (Molecular Machines & Industries, #50102) membrane slides, then stained with haematoxylin & eosin (H&E) using a Leica XL automated tissue stainer. Sections mounted onto glass slides were evaluated by a clinical pathologist and regions of ‘tumour core’ and ‘leading edge’ were annotated to guide dissection. From sections mounted on MMI slides, equal volumes (0.05mm^3^) of tumour core and leading edge were dissected from multiple sections using a MMI CellCut Laser Microdissection system. Tissue was collected using MMI transparent isolation caps and pooled with dissected tissue from the same region.

### Protein extraction from FFPE tissue

Formalin-mediated protein cross linking was reversed by resuspending the dissected tissue in 50mM TEAB (triethyl ammonium bicarbonate) containing 5% SDS (w/v) and heating at 95°C for 20mins, then 60°C for 2 hours. To assist the solubilisation of extracellular matrix (ECM) proteins, urea and DTT (dithiothreitol) was added to samples to a final concentration of 8M and 5mM respectively. Samples were then sonicated in a LE220-Plus focused ultrasonicator (Covaris) for 10 mins.

### Protein extraction from snap frozen tissue

Approximately 1 mm^3^ of snap frozen tissue was resuspended in 5% SDS (w/v), 8M urea, 5mM DTT, 25mM ammonium bicarbonate, cOmplete mini EDTA-free protease inhibitor cocktail (pH7). Resuspended tissue was homogenised via sonication in a ‘LE220-Plus’ focused ultrasonicator (Covaris) for 10 mins.

### Sample preparation for mass spectrometry (MS) proteomics

Either 50 μg protein (snap frozen tissue) or all extracted protein (FFPE tissue) was reduced in 5% (w/v) SDS with 2.5 mM DTT, incubated at 60 °C for 10 min. Iodoacetamide (IAA) was then added to a final concentration of 15 mM, vortexed and incubated in darkness at room temperature for 30 min. IAA was quenched through addition of DTT and the solution cleared by centrifugation for 10 min at 14000 RCF (relative centrifugal field). Samples were acidified by addition of phosphoric acid to 1.2% (w/v), and their volume increased six-fold with the addition of binding buffer (90% methanol in 100 mM TEAB, pH 7.1). Samples were then immobilised and digested on S-trap spin columns (ProtiFi), according to the manufacturer’s instructions. Resulting peptides were desalted, in accordance with the manufacturer’s protocol, using POROS R3 beads (Thermo Fisher) and lyophilized using a SpeedVac (Thermo Fisher scientific).

### Liquid chromatography coupled tandem mass spectrometry (LC-MS/MS)

Dried peptides were resuspended in 10 μL 0.1% formic acid in 5% acetonitrile (ACN). Samples were analysed using an ultiMate^®^ 3000 Rapid Separation LC system (RSLC, Dionex Corporation) coupled to a Q Exactive HF Mass Spectrometer (Thermo Fisher). The mobile phase A was 0.1% formic acid in water and B was 0.1% formic acid in ACN. The solvent gradient was adjusted: 95% A and 5% B to 18% B at 58 min, 27% at 72 min, and 60% at 74 min, using a flow rate of 300 nL/min and a 75 mm x 250 μm inner diameter CSH C18 analytical column (Waters). Peptides were automatically selected for fragmentation by data dependent acquisition (DDA); data was acquired for 60 min in positive mode.

### Proteomics Data Processing

Raw mass data were processed using MaxQuant (v1.6.14.0, available from Max Planck Institute of Biochemistry) (35). Features were identified using default parameters in MaxQuant, then searched against the human proteome (UniProt database, August 2020). Oxidation of methionine (M) & proline (P) and Acetylation of protein N-terminus were set as variable modifications. Carbamidomethyl (C) was set as a fixed modification. Peptide quantitation was performed using label-free quantification (LFQ), using only unmodified, unique peptides and with ‘match between runs’ enabled (unless explicitly stated otherwise within the main text).

Statistical analysis was performed using MSqRob (36, 37). LFQ data was normalised using the median of peptide intensities. Sample region (‘core’ or ‘edge’) was treated as a fixed effect. Run, peptide sequence and replicate (biological or technical, experiment depending) were treated as random effects. Peptides belonging to contaminant protein lists (as annotated by MaxQuant) or proteins with few that two peptides were excluded from statistical analysis.

### Ontological analysis of proteomics data

Annotation of matrisome proteins was achieved by screening all identified proteins against MatrisomeDB, a curated database of ECM proteins (38–40). Pathway analysis was performed on differential protein abundance data generated by MSqROB using PANTHER’s (Protein Analysis Through Evolutionary Relationships) ‘statistical enrichment’ test (2019 v.16, released Feb-21) (41). False discovery rate (FDR) correction was applied, and proteins were grouped by ‘Reactome pathways’ (41,42).

### Immunohistochemistry (IHC)

IHC was performed as detailed in Herrera et Al., 2019 (43). In brief, sections were subjected to antigen retrieval using citrate buffer, pH6.0 (BioCare, RV1000) for 30mins at 100°C. Sections were then stained with anti-TNR rabbit polyclonal (AbCam, #ab198863) or anti-FN1 rabbit polyclonal (AbCam, #ab23750) and staining visualised using a Novolink Polymer Detection Systems (Leica, RE7200-CE) as per manufacturer’s instructions. Images were acquired on a 3D-Histech Pannoramic-250 microscope slide-scanner using a 20x/ 0.80 Plan Apochromat objective (Zeiss). Snapshots of the slide-scans were taken using the Case Viewer software (v2.5; 3D-Histech). Quantification of staining was performed using the QuantCenter (v2.0) plugin for CaseViewer. In brief, a mask was drawn around each region of interest selected by a clinical pathologist. For each section, a second mask was generated with RBG thresholds set to selectively captured only stained tissue. The proportion of the stained area within each region was then calculated as a percentage relative to the total area.

## Results

### Laser-capture microdissection coupled with mass spectrometry allows for spatially resolved proteomic analysis of FFPE glioblastoma tissues

To visualise the histological features of tissues during LCM, tissue samples must be sectioned, stained, and mounted onto microscope slides. Formalin-fixation, paraffin-embedding (FFPE) and paraffin microtomy is the standard methodology for processing tissues for diagnosis. However, formalin fixation generates crosslinks between amino-acid residues which may adversely impact the ability to isolate, detect and quantify protein species via LC-MS/MS. Thus, we compared the global and ECM-specific proteomic landscapes identified from laser microdissected FFPE glioblastoma tissue and equatable snap-frozen tissue.

To avoid confounding issues from genetic variation, tumour site or surgical procedure etc., comparable snap-frozen and FFPE tissue samples were generated by bisecting a single biopsy specimen into two fragments of similar volume that were then processed independently, [Fig.1.a.]. From the FFPE tissue, 0.05 mm^3^ regions of tumour core and infiltrating edge were excised and collected by LCM from serial tissue sections, [Fig.1.b.]. In parallel, a 1mm^3^ fragment of the contiguous frozen tissue was isolated and homogenised. The dissected FFPE and frozen tissues were trypsinised and desalted contemporaneously, then analysed by LC-MS/MS. The raw mass spectrometry data from each sample were treated independently during computational analysis, ensuring each protein identification was established by secondary peptide fragmentation, rather than through similarities between successive samples.

**Figure 1.**
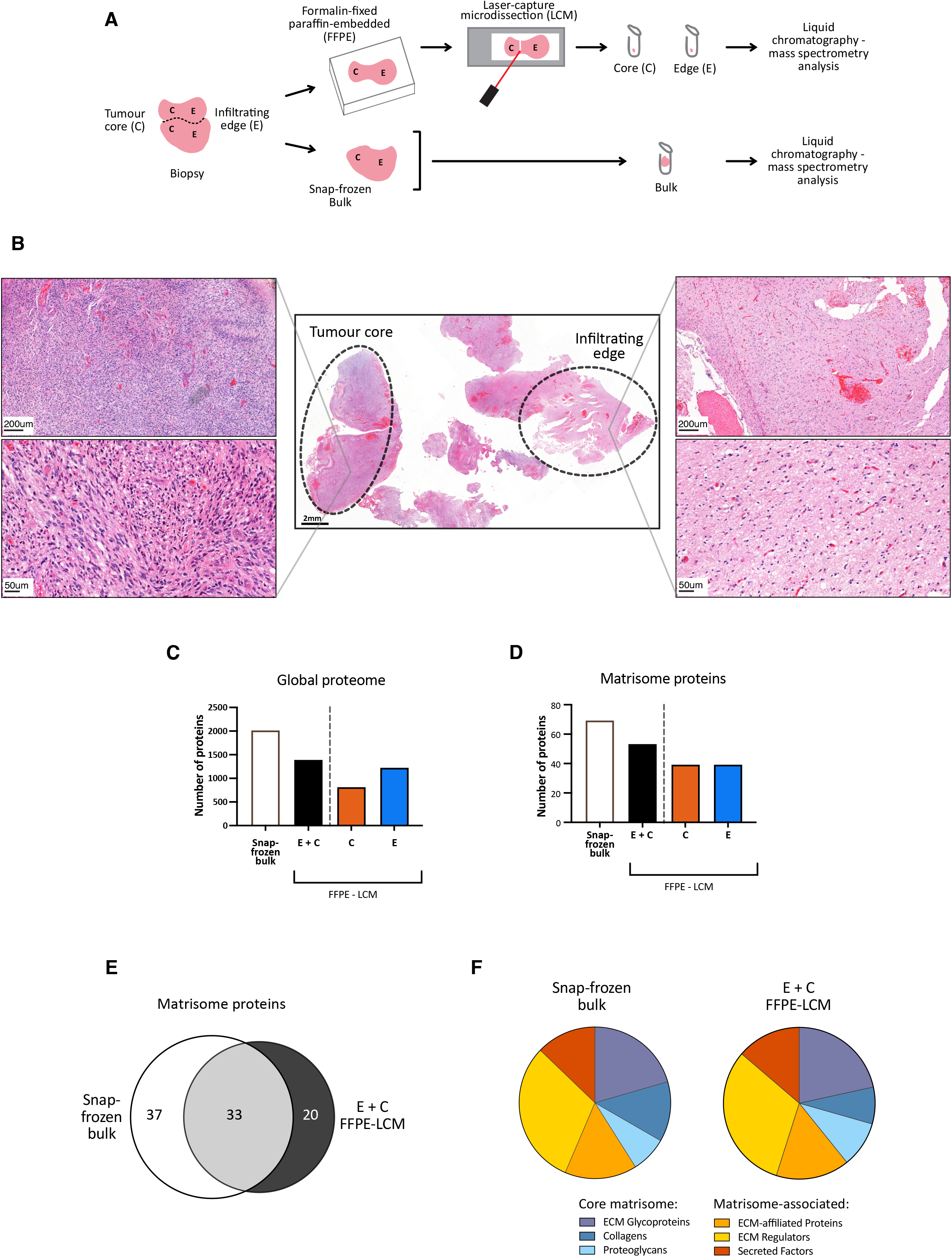
Proteomic analyses of glioma tissues, comparing snap-frozen, bulk samples vs. formalin-fixed, paraffin-embedded (FFPE) and sectioned samples. (**A**) Schematic representation of sample preparation for mass spectrometry (MS) proteomics: brain tumour biopsies were either sectioned using laser-capture micro-dissection (LCM) into core (C) and infiltrating edge (E) regions, following FFPE treatment, or used whole, following snap freezing in liquid nitrogen. (**B**) An FFPE section of tumour tissue, with haematoxylin and eosin (H&E) stain, and annotation of tumour core (C) and the infiltrating edge (E) regions. (**C**) Number of proteins identified by MS in bulk analysis of brain tumour tissue, in LCM-coupled MS analysis of core (C), edge (E), and in the combined set of core and edge (C+E). (**D**) The same analysis, but considering only matrisome proteins, as defined by Naba et al (2012)(38). (**E**) Venn diagram showing overlap of matrisome proteins in MS analysis of snap-frozen bulk tissue vs. proteins detected in the combined set of core and edge (C+E) regions. (**F**) Comparison of the matrisome categories detected by snap-frozen bulk analysis vs. LCM (38).

As the area sampled from the snap-frozen tissue was selected without any segregation of the core or edge regions (i.e. represents the ‘bulk’ tissue), we initially compared the protein species identified by MS analysis of the frozen tissue to the combined list of proteins from both LCM tissue sites (E + C). In total, 2007 proteins were identified in the frozen tissue, while 1384 proteins were identified in the combined E + C LCM sample, with 1218 and 808 proteins in the edge and core samples respectively [Fig.1.c.]. ‘Core ECM’ and ‘ECM-associated’ proteins can be grouped together into several hierarchical categories based on their function and structural properties that together describe the ‘Matrisome’ (38–40). 69 Matrisome proteins were identified in frozen tissue, compared with 53 in the LCM combined edge and core set (39 matrisome proteins were observed in both core and edge samples), [Fig.1.d.]. Approximately half of the matrisome proteins identified in the frozen bulk tissue were also identified in the LCM samples, [Fig.1. e.]. Furthermore, analyses of frozen and LCM samples contained comparable proportions of proteins for each of the matrisome subcategories (core matrisome: ECM glycoproteins, collagens and proteoglycans; matrisome-associated: ECM-affiliated proteins, ECM regulators and secreted factors) [Fig.1.f.]. These results indicate that our methods reverse formalin-induced cross-linking in FFPE sections of brain tissue and can enable spatially resolved proteomic analysis, with a coverage of matrisome proteins comparable to that achieved with bulk, snap-frozen tissues.

### LCM-proteomic analysis allows spatially resolved and reproducible interrogation of the glioblastoma microenvironment

We next assessed the reproducibility of LCM-proteomics from FFPE tissue. In triplicate, regions of tumour core and edge were dissected and analysed from serial sections of a single specimen [Fig.2.a.]. Variation in protein abundances between the technical replicates and regions were then visualised using principal-component analysis (PCA), and by computing Pearson’s correlation coefficients for pairwise comparisons of protein abundance [Fig.2.b,c.]. The PCA showed clear clustering of core and edge samples in the primary principal component axis (PC1) accounting for 62.5% of the variance, indicating that variance between intra-tumour regions was greater than that between technical replicates. This finding was confirmed by a comparison of Pearson’s correlation coefficients, where R^2^ values ranged from 0.80 – 0.97 for comparisons of replicate samples within like regions, and between 0.44 – 0.56 for samples from different regions, [Fig.2.c.].

**Figure 2.**
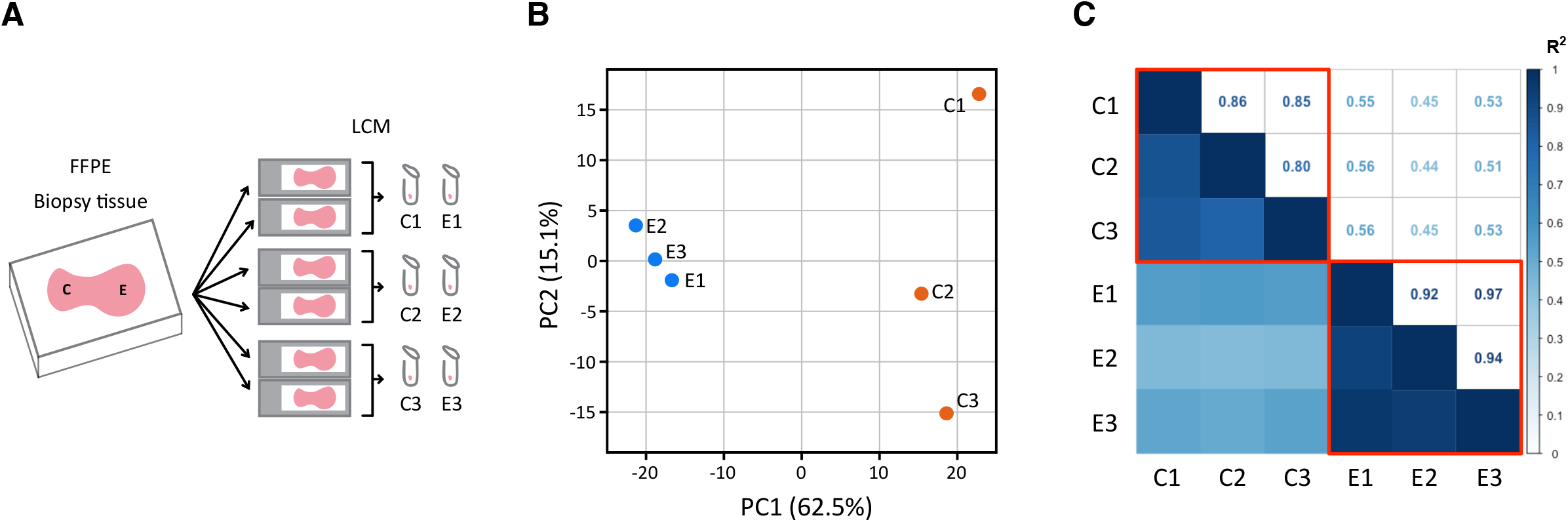
Laser-capture micro-dissection (LCM) coupled mass spectrometry (MS) analysis of formalin-fixed, paraffin-embedded (FFPE) brain tissue is reproducible. (**A**) Schematic representation of the experiment: tumour core (C) and infiltrating edge (E) regions were taken from consecutive sections through an FFPE tissue sample, providing a source of spatially-matched replicates. (**B**) Principle component analysis of peptide-level MS data, showing separation of core and edge regions in the axis with highest variance (principal component 1, PC1). (**C**) Matrix of Pearson’s R-squared correlations between MS datasets.

Both PCA and analysis of Pearson’s R^2^ identified variance to be greater between replicates taken from within the tumour core (spread of points in secondary principal component axis, PC2; R^2^ range, 0.80 to 0.86) than within the edge (clustering in both PC1 and PC2; R^2^ range, 0.92 to 0.97). This result indicates greater heterogeneity within the tumour core microenvironment than at the infiltrating tumour edge [Fig.1.b.].

### LCM-proteomics identifies distinct extracellular matrix proteomic signatures associated with the glioblastoma tumour core and infiltrating edge

Having established the reproducibility of LCM-proteomics to examine both the global and ECM-specific proteomic landscapes of FFPE glioblastoma tissue, we next employed the technique to interrogate differences between the microenvironments of the tumour core and infiltrating edge. Regions of core and edge were laser-microdissected from three independent tumours and analysed via LC-MS/MS. 1107 proteins were identified in total, with 79 proteins showing significant differential abundances between the core and edge regions [Fig.3.a.].

**Figure 3.**
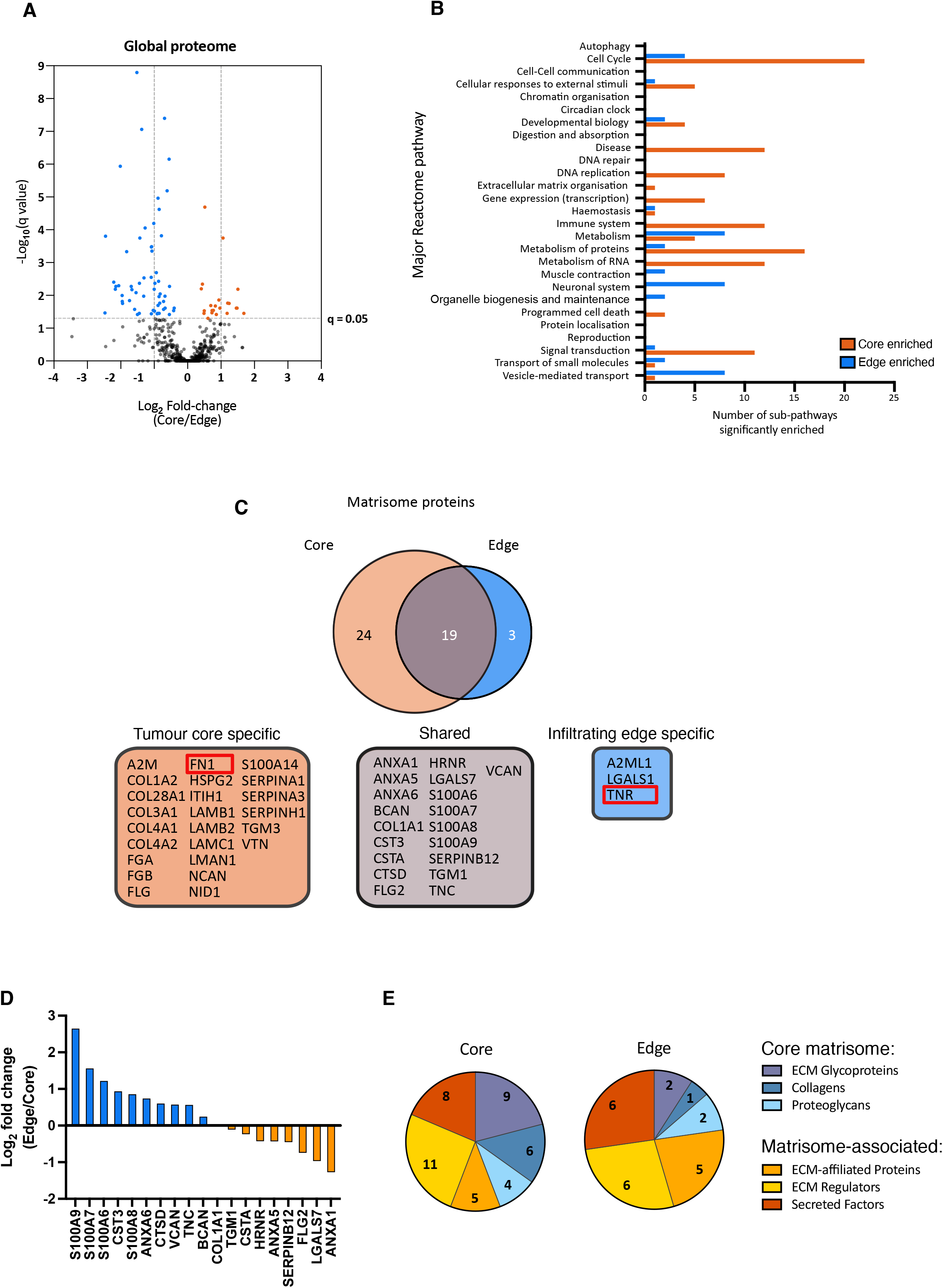
Quantitative comparison of core and infiltrating edge sections of brain tumour tissue reveals characteristic protein composition. (**A**) Volcano plot showing analysis of infiltrating edge vs. tumour core tissue samples, analysed by mass spectrometry (MS) proteomics following formalin-fixation, paraffin embedding and sectioning by laser-capture micro-dissection (Figure 1A). Proteins significantly enriched in core and edge are shown in orange and blue, respectively (n = 3 biological replicates). (**B**) Plot showing the number of Reactome pathways significantly enriched (false-discovery corrected p-value (q val) < 0.05) in the highest event hierarchy, comparing infiltrating core to tumour edge samples (41,42).(**C**) Venn diagram showing matrisome proteins identified with two-or-more unique peptides in the tumour core and infiltrating edge sections. This analysis identified proteins with specificity to different regions of the tumour, for example, fibronectin (FN1) in the core, and tenascin-R (TNR) in the edge. (**D**) Abundance fold change of Proteins identified in both core and edge condition, (Figure 3C). Proteins enriched in core/edge are shown in orange or blue respectively (n = 3 biological replicates). (**E**)Analysis of core and edge matrisome proteins by category (38). This analysis suggested lower levels of structural matrisome components such as collagens in the edge vs. the core.

To examine the differences in the biological processes between the cores and leading edges of glioblastoma, pathway analysis was performed on the differential protein abundance dataset [Fig.3.b.], [Sup. Table 1]. Pathways associated with cellular proliferation, including ‘cell cycle’, ‘DNA replication’, ‘gene expression’, ‘metabolism of RNA’, and ‘metabolism of proteins’, were more commonly enriched in the tumour core compared with the leading edge. This was consistent with prior reports that the tumour core is residence to a population of rapidly proliferating cancer cells. Furthermore, pathways related to the ‘immune system’ were only found to be enriched in core tissue and not in the infiltrating edge, consistent with this region being pro-inflammatory with extensive infiltration of immune cells (44, 45). Finally, multiple pathways associated with ‘neuronal systems’ were significantly enriched in the tumour edge, while these were absent in the core. This suggested that the infiltrating edge retained some degree of normal neural architecture.

We next asked whether the tumour core and infiltrating edge each had a characteristic ECM composition. MS data from the tumour core and edge were reprocessed independently to identify the protein species present in each region, and the resultant protein lists screened for matrisome proteins [Fig.3.c.]. 43 and 21 matrisome proteins were identified in the core and edge respectively. Of these components, 24 matrisome proteins were uniquely present in the core, and 3 exclusively in the invasive edge. 19 matrisome proteins were found to be common to both regions but showed differences in relative abundances across them [Fig.3.d.]. A comparison of the matrisome proteins subclasses observed in the two regions revealed that the tumour core was enriched for structural matrisome components, including ECM glycoproteins, collagens and proteoglycans, relative to the infiltrating edge tissue. In contrast, the infiltrating edge was predominantly composed matrisome-associated proteins [Fig.3.e.]. This is in agreement with observations that the core tumour microenvironment becomes increasingly ECM dense and mechanically stiffer during disease progression(9).

We next sought to independently validate these specific ECM signatures with immunohistochemistry. Fibronectin and tenascin R were selected as they were detected by LC-MS/MS exclusively in either the core or edge respectively. Immunostaining of 5 glioblastomas with a fibronectin-specific antibody confirmed it was heterogeneously distributed across the tissues [Fig.4.a.]. Threshold-based image quantification demonstrated that a greater relative area of tumour core displayed positive fibronectin staining than the infiltrating edge in four out of five tumours examined, consistent with the mass spectrometry data from this region [Fig.4.b.]. In contrast, immunostaining for tenascin-R (TNR) showed protein expression to be higher at the leading edge than the tumour core across all five tumours examined, with a median protein density ~50% lower in the core than the invasive edge [Fig.5.a,b.]. Two of the five specimens contained regions of normal brain parenchyma. For these normal tissue regions, anti-TNR antibody staining was greater than that present at the tumour edge, suggestive of a gradient in the abundance of TNR between the normal brain parenchyma, the glioblastoma infiltrating edge and the tumour core [Fig.Ex.5.a,b.].

**Figure 4.**
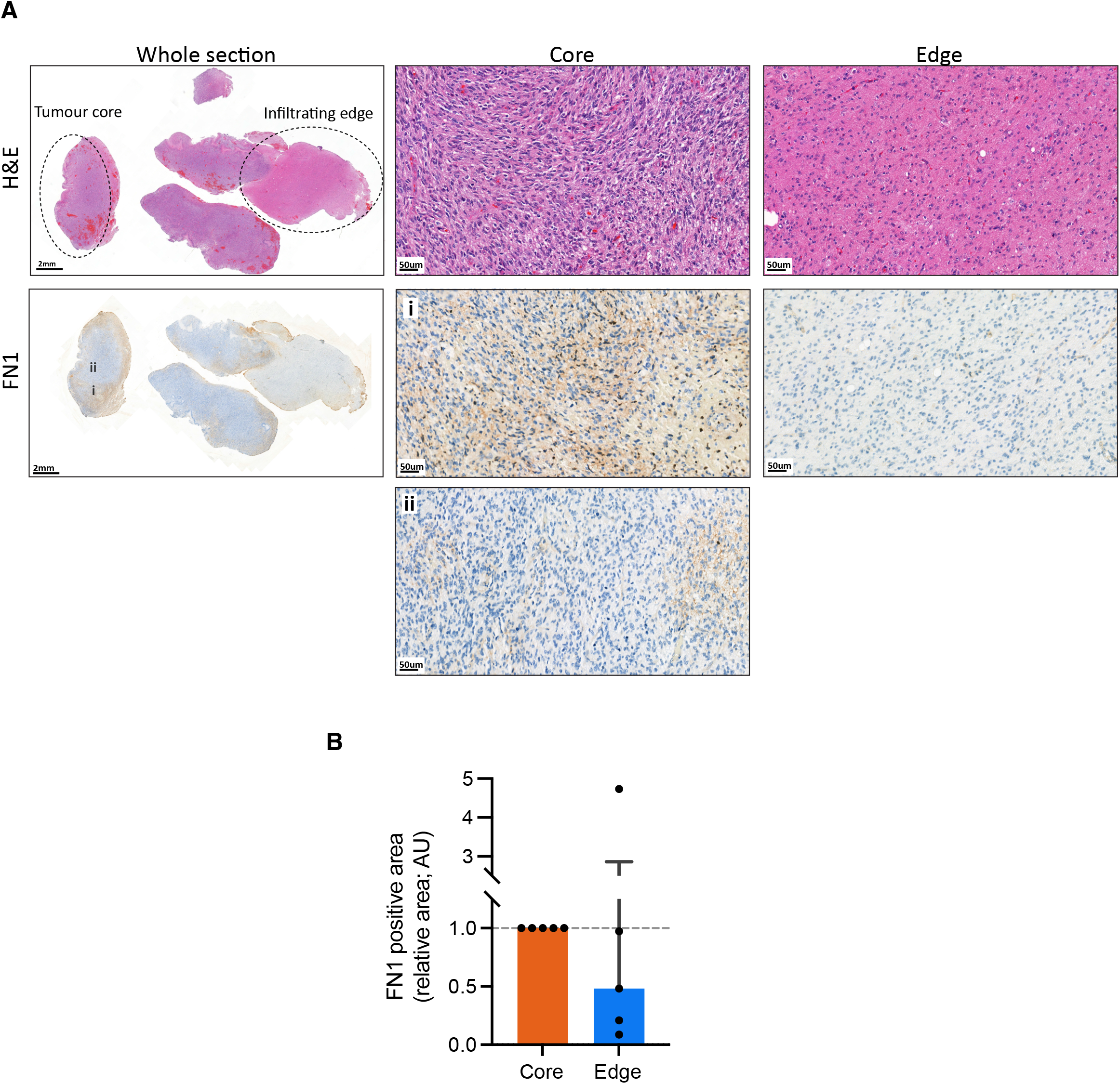
Immunohistological staining of fibronectin (FN1) shows variability within tumour core. (**A**) A section of glioblastoma tumour, with annotation of core and infiltrating edge, with haematoxylin, eosin (H&E) and anti-FN1 staining. Variation in FN1 stain was identified within the tumour core region, exemplified by the regions annotated (i) and (ii). (**B**) Quantification of relative regional areas with positive FN1 staining for tumour core and edge (normalised to specimen-matched core staining, n = 5 biological replicates). Median and interquartile ranges are displayed

**Figure 5.**
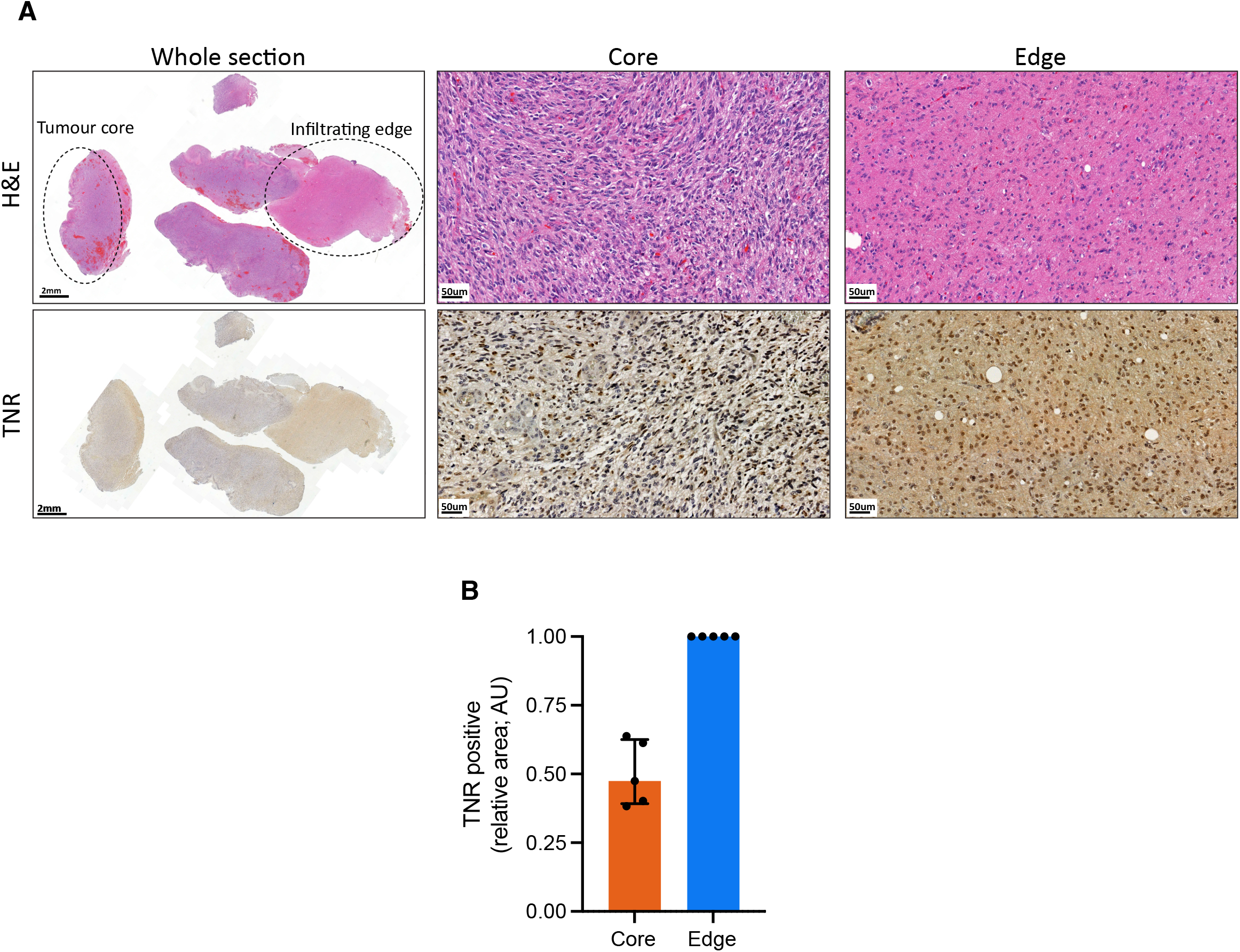
Immunohistological staining shows reduced levels of tenascin-R (TNR) at the core of the tumour, relative to the Infiltrating edge. (**A**) A section of glioblastoma tumour, with annotation of core and infiltrating edge, with haematoxylin, eosin (H&E) and anti-TNR staining. (**B**) Quantification of relative regional areas with positive TNR staining for tumour core and edge (normalised to specimen-matched edge staining, n = 5 biological replicates); the infiltrating edge of the tumour showed consistent enrichment for TNR. Median and interquartile ranges are displayed

Together these findings confirmed the results of our LCM-proteomics and demonstrated that this approach is capable of spatially resolved interrogation of the regional extracellular matrix microenvironment of glioblastoma.

## Discussion

Glioblastoma is characterised by vast divergence between the microenvironments of the tumour core and the infiltrating edge. Standard-of-care therapy involves maximally safe surgical resection, typically eliminating the majority of the core tissue. However, 90% of recurrences occur from the residual infiltrative edge at the resection boundary. Despite this evidence, studies investigating glioblastoma ECM composition have focused on the tumour core, and the composition of the infiltrating edge is often overlooked (1,3, 10, 33). Here, we demonstrated that LCM-proteomics could compare the TME of the core and infiltrative edge of glioblastoma, identifying characteristic ECM signatures for each of these two compartments. To our knowledge, this is the first time LCM-proteomic analysis has been applied to primary human glioblastoma tissues, and has advanced our understanding of the heterogeneous glioblastoma microenvironment.

LCM-proteomics allows spatially precise and technically reproducible exploration of the proteomic landscape of human tissue microenvironments. Previously, LCM-proteomic analysis has been applied to frozen tissues from patient-derived xenograph (PDX) glioblastoma models (46). However, PDX models often lack many of the TME constituents present in the primary tumour, including the native human ECM, vasculature, and non-neoplastic cell populations, and thus, may only recapitulate the cellular features of human tumours. To interrogate the native ECM of glioblastoma, we applied LCM-proteomics to FFPE tissue biopsied directly from primary tumour sites. Patient derived FFPE tissue for LCM-proteomics offers several practical advantages. FFPE biopsies are routinely generated as part of the clinical pathology pipeline and can be archived for years without detriment. Archived FFPE materials stored in pathology archives represent a sizeable and readily available source of clinically annotated tissues for retrospective studies on the glioblastoma TME. We demonstrate here that formalin-fixation and paraffin-embedding does not impede ECM protein identification or quantitation by LCM-proteomics. Therefore, the data derived from regional niches in our study can be considered representative of the *in vivo* TME and spatial variation in the glioblastoma ECM landscape.

Our spatial proteomic analysis demonstrated that the glioblastoma core consisted of a more complex milieu of matrisome components than the infiltrating edge. Fibrillar collagens collagen alpha-2 (I) chain and collagen alpha-1 (III) chain (COL1A2 and COL3A1 respectively) were exclusively identified in the tumour core, whereas collagen alpha-1(I) chain (COL1A1) was identified in both core and edge. Furthermore, pathway analysis highlighted collagen biosynthesis and remodelling as being significantly enriched in the tumour core relative to the leading edge [Sup. Table 1]. Fibrillar collagens are expressed at very low levels in normal adult brain parenchyma, contributing to the organ’s soft mechanical properties (47). The expression of fibrillar collagens in glioblastoma has been a point of contention in the field, with some studies reporting their absence within the tumour mass (15, 48). However, the data presented here, along with evidence from recent transcriptomic datasets and IHC studies (12, 13, 49), together prove the presence of fibrillar collagens within the tumour stroma, and partially explains the mechanically stiffer and more fibrous microenvironment of glioblastoma tissues relative to normal brain parenchyma.

The tumour core was also enriched with proteins classically associated with both the normal brain parenchyma and the basement lamina, including collagen IV (COL4A1, COL4A2), laminin I (LAMB1, LAMC1), and perlecan (HSPG2), relative to the infiltrating edge (50). These data correlate with the previously reported findings that ECM components of the normal brain parenchyma become dysregulated during glioblastoma instigation and disease progression (14–20), and/or, with the increased presence of vascular structures within the tumour core due to microvascular hyperplasia, a defining histological feature of the tumour core.

Fibronectin (FN1) was detected exclusively in the tumour core by LCM-proteomics, while IHC revealed that, for four out of five cases examined, the average expression of fibronectin across the tumour core was greater than at the leading edge. Fibronectin is typically absent from the parenchyma of the normal neocortex, but is instead associated with the residing vasculature, in the form of insoluble fibrils associated with the basement membrane and as a soluble component of plasma (51, 52). Fibronectin specific staining of our patient cohort demonstrated that distribution of fibronectin was heterogeneous across tumour core regions, with strongest staining typically observed in areas proximal to thrombi, suggesting expression may be associated with ECM remodelling in response to ‘wound-like’ damage occurring within the core (53).

Few proteins were identified exclusively in the infiltrating edge with LCM-proteomics. However, relative quantitation of matrisome proteins shared between the core and the infiltrating edge indicated that a number show differential expression between the two tumour compartments. These include versican (VCAN) and tenascin-C (TNC), both showing increased expression in the invading edge. Tenascin-C is typically expressed at low levels in normal brain, but has been implicated in glioma, where high expression has been linked to recurrence (54). Versican is a member of the lectican family of chondroitin sulfate proteoglycans (CSPGs). Other family members identified in our analysis include brevican (BCAN) and neurocan (NCAN), the latter only detected in the tumour core. The lecticans have been implicated in the function of the perineuronal net (PNN), a specialised ECM surrounding certain populations of neurones. Other components of the PNN include tenascin-R (55). Tenascin-R was detected by LCM-proteomics exclusively in the invading edge, which may reflect its association with PNNs in normal tissue, into which the tumour edge is invading. Supporting this, IHC data from sections containing regions of tumour core, edge and normal tissues suggest that a gradient of tenascin-R abundance may exist between the core and adjacent normal tissue, with tenascin-R abundance lowest in the central core and highest in the non-transformed parenchyma. Galectin 1 (LGALS1) was also exclusively detected in the invading edge, and its expression has been linked to poor overall survival (56).

Here we demonstrate that LCM-proteomics offers a viable methodology for exploring the composition of distinct histological regions, and thus tumour heterogeneity, of glioblastoma in archived clinical samples. However, even within the area we describe herein as ‘tumour core’, smaller scale, discrete, microanatomical features exist, such as pseudopalisading necrosis and microvascular hyperplasia, each of which have unique cellular and non-cellular microenvironments that contribute to the pathobiology of glioblastoma (57, 58). The spatial resolution of laser-microdissection is around 3 μm, sufficient to allow the isolation of these microanatomical regions (32, 59, 60). Indeed, a recent study utilised laser-capture microdissection coupled to RNA-sequencing to isolate and transcriptionally characterise six distinct histological compartments within glioblastoma (33). An exciting next step will be combining LCM-proteomic and LCM-transcriptomic analyses using serial tissue sections for a comprehensive understanding of the cellular and non-cellular constituents of the distinct regional niches of glioblastoma.

## Personal Acknowledgements

Our thanks goes to David Knight, Stacey Warwood, Ronan O’Cualain and Julian Selley of the UoM Biological Mass Spectrometry Core Facility for their assistance running the HPLC and mass spectrometry systems. We are also particularly grateful to Peter March and Roger Meadows of the UoM Bioimaging facility their help with the microscopy. We thank Craig Lawless for his assistance with bioinformatics and data analysis; and Jeremy Herrera for his advice on the LCM-proteomics methodology. We also thank the staff of the department of Neurosurgery, Manchester Centre of Clinical Neuroscience, in particular Helen Mayers for her assistance in staining tissues.

## Author contributions

Investigation - RP, DP, LD & JM; Formal Analysis, RP and EA; Clinical sample acquisition – PIDU & ID; Writing – Original Draft, RP; Writing – Visualization, Review & Editing, RP, DP, LD, EA, AG, FR and JS; Project Administration and Funding Acquisition, AG, FR and JS.

