## Extended Figure 1 for "Novel Laser Capture Microdissection-Proteomic Analysis Identifies Spatially Distinct Extracellular Matrix Signatures in the Core and Infiltrating Edge of Human Glioblastoma"

Ex. 5

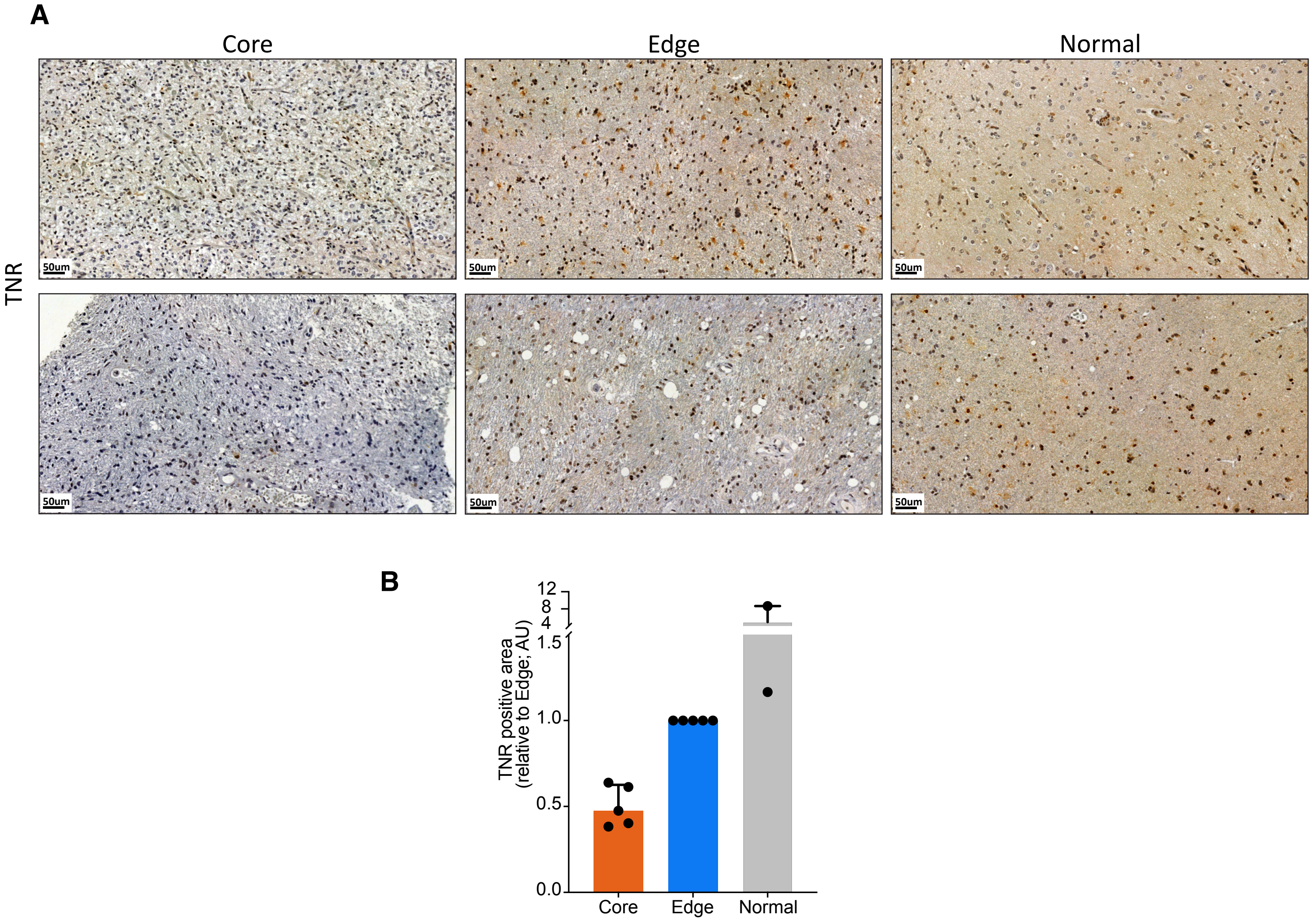


Extended figure 5.

**Immunohistological staining shows reduced levels of tenascin-R (TNR) at the core of the tumour, relative to the Infiltrating edge and normal parenchyma** (**A**) Section of human glioblastoma tumour, with annotation of core, infiltrating edge and normal tissue, with haematoxylin & eosin (H&E) and anti-FN1 staining. (**B**) Quantification of relative areas of regions with positive TNR staining in tumour core, infiltrating edge and normal tissue (normalised to specimen-matched edge staining, n = 2 biological replicates).
